# Single-cell transcriptome landscape of zebrafish liver reveals hepatocytes and immune cell interactions in understanding nonalcoholic fatty liver disease

**DOI:** 10.1101/2022.02.06.479276

**Authors:** Yingyi Huang, Xiang Liu, Hong-Yan Wang, Jian-Yang Chen, Xianghui Zhang, Yubang Li, Yifang Lu, Zhongdian Dong, Kaiqiang Liu, Zhongduo Wang, Qian Wang, Guangyi Fan, Jun Zou, Shanshan Liu, Changwei Shao

## Abstract

Zebrafish have emerged as an attractive animal model for studying nonalcoholic fatty liver disease (NAFLD). However, little is known about the cell types and intercellular interactions in zebrafish liver. Here, we established a liver atlas that consists of 10 cell types using single-cell RNA sequencing. By examining the heterogeneity of hepatocytes and analyzing the expression of NAFLD-associated genes in the specific cluster, we provide a potential target cell model to study NAFLD. Additionally, our analysis identified two distinct resident macrophages with inflammatory and noninflammatory functions and characterized the successive stepwise development of T cell subtypes in the liver. Importantly, we uncovered possible molecular mechanisms and revealed the central regulation of macrophages on target cells of fatty liver by analyzing the cellular interaction between hepatocytes and immune cells. Our data provide valuable information for future research on NAFLD in zebrafish.

## Introduction

The liver is a vital organ in the body of vertebrates that exerts essential metabolism and immune function (1). Liver dysfunction leads to a series of diseases, including viral hepatitis, tumors, fibrosis, cirrhosis and fatty liver (2). Among them, nonalcoholic fatty liver disease (NAFLD) has become a common cause of chronic liver disease worldwide, accounting for 24% of liver diseases (3). Typical NAFLDs include nonalcoholic fatty liver (NAFL), nonalcoholic steatohepatitis (NASH), liver fibrosis, liver cirrhosis and NAFLD-related liver cancer. The progression of NAFLD to liver fibrosis and cirrhosis greatly increases the mortality of liver-related diseases. However, the pathogenesis of NAFLD has yet to be fully elucidated due to the complexity of mechanisms and the ethical issues involved in human experiments. Thus, the establishment of a NAFLD animal model in which the mechanism of liver function has been relatively well studied is of great clinical importance.

Currently, the “multiple hit” hypothesis has been widely accepted to explain the complexity of NAFLD (4). This concept involves insulin receptor (IR), lipotoxicity, inflammatory response, genetic polymorphism and epigenetics, adipokines and liver factors, bile acid (BA), and gut microbiota (GM) (5). The accumulation of lipid toxic substances caused by the impairment of hepatocyte lipid metabolism is the key to the occurrence of nonalcoholic steatohepatitis (NASH) (6). Other factors, such as dysfunction of cytokines and adipokines, ATP deficiency, uric acid toxicity and products of intestinal microorganisms, may also be involved in regulating the development of lipotoxic stress, injury and inflammation of hepatocytes (7).

Among the many factors affecting the occurrence and development of NAFLD, the important role of immune cell activation should not be ignored (8). Abnormalities in the liver systemic immune response were found to be important factors in the progression of NAFL to NASH and liver fibrosis (9). Macrophages are involved in the pathogenesis of NAFLD (10-12). In terms of cellular functions, liver macrophages can be divided into M1 (inflammatory) and M2 (noninflammatory) macrophages. M1 macrophages promote the production of inflammatory cytokines and reactive oxygen species and are involved in the progression of NAFLD liver inflammation. In contrast, M2 macrophages execute immune regulation, inhibit parasitic infection and strengthen tissue remodeling and are involved in the progression of NAFLD liver fibrosis (13-15). Macrophage-associated factors such as pattern recognition receptors, complement molecules and STING are suggested to promote the development of NAFLD. In addition, T cells are critical in maintaining liver immune homeostasis and restricting the progression of liver diseases (16). CD4^+^CD25^+^FOXP3^+^ T cells (regulatory T cells, Tregs) play an important role in blocking the transformation of NAFL into NASH and mediating the remission of NASH inflammation and fibrosis (17-19). Additionally, excessive production of Th1 cell-derived proinflammatory cytokines may promote hepatic insulin resistance and NASH (8). A recent study has shown that CD8^+^ T cells undergo autoaggression through a series of continuous activation steps to mediate the occurrence of liver cancer in NASH patients (20, 21).

Zebrafish (*Danio rerio*) has become an important model to study liver development and diseases and understand the pathogenesis of major liver diseases. Zebrafish and mammals are similar in hepatocyte composition, function and genetics. However, our understanding of hepatocyte biology in zebrafish models has largely come from knowledge of hepatocytes in juvenile fish obtained using image tracking approaches when the adaptive immune system is not well developed (*22*), which limits the in-depth study of liver fibrosis and liver cirrhosis. To date, the landscape of human and mouse liver cells has been explored at single-cell resolution, providing valuable information to elucidate liver functions and diseases (*23-26*). The single-cell atlas of adult zebrafish liver is of great significance to complement mammalian studies to uncover the pathogenesis of liver diseases from evolutionary perspectives.

In this study, we used high-throughput scRNA-seq to undertake an unbiased examination of the cellular landscape of healthy zebrafish liver. We identified 10 cell types using known lineage marker genes and potentially targeted cell models for hepatocytes by analyzing cell subclusters. Our analysis revealed remarkably heterogeneous transcriptomic signatures for the pro- and noninflammatory macrophage subtypes and that associated with the development of T cell subtypes. Importantly, comparative analysis of unbiased cell interactions between zebrafish and human orthologous genes uncovered possible molecular mechanisms of the interactions between immune cells and hepatocytes in zebrafish liver and the roles of macrophages in regulating target cells of fatty liver. Our data provide valuable information for future investigation on NAFLD in zebrafish and enhance our understanding of the molecular events and cellular mechanisms related to liver functions and diseases.

## Results

### Single-cell atlas of zebrafish livers

To investigate the diversity of zebrafish liver cell types, zebrafish liver tissue cells were prepared for preparation of single-cell libraries using the MGI2000 platform (Figure 1A). Six single-cell libraries were sequenced at a median of 78,178 reads per cell, which almost reached saturation, as indicated by read down sampling analysis. Subsequently, after quality filtering, a total of 4268 cells were retained, with a median of 4941 transcripts generated per cell. The Seurat package was used for single-cell data analysis and batch effect removal. Cells were further filtered to reduce the dimensionality of the cells, and 10 clusters were obtained (Figure 1B and 1C). Comparing the proportions of different samples in the same cluster, we found that in each independent sample, the proportion of cells in the same cluster was similar, indicating that our data were unaffected by batch effects and could be used to interpret biological significance (Figure 1D).

**Figure 1.**
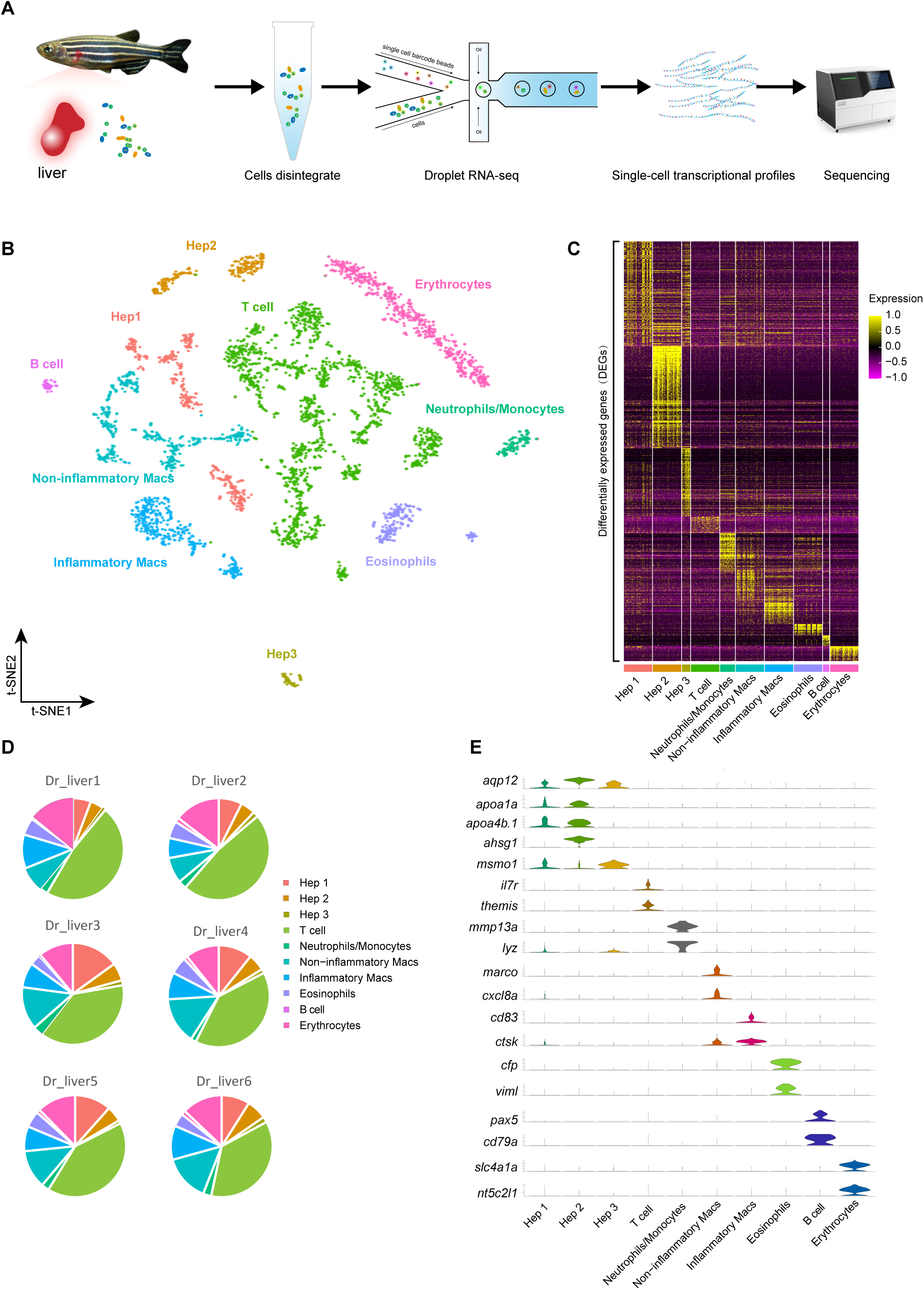
Overview of 10 cell types determined by scRNA sequencing in zebrafish. **A**. Schematic representation of the single-cell RNA-seq analysis of zebrafish liver cells. **B**. Visualization of major testis cell types for 4268 cells in UMAP (unknown is undefined). Different cell types are shown in distinct colors. **C**. Heatmap showing the expression of the DEGs for the main cell types. Z-scores were calculated by subtracting the average value for the set of data from the value for each cell and dividing by the standard deviation. **D**. Pie chart showing proportion of 10 cell types. E. Violin plots of the normalized expression of marker genes for 10 cell types.

Based on T-distributed stochastic neighbor embedding (t-SNE) analysis and differentially expressed gene analysis, we annotated three nonimmune populations and seven immune cell populations. Nonimmune cell types primarily consisted of three hepatocyte subclusters: Hepatocyte (Hep) 1 expressing *aqp12* (*27*) and *itih2* (*28*), Hep 2 expressing *ahsg1* (*29*), *apoa1a* (*30*) and *apoa4b*.*1* (*31*), and Hep3 expressing *msmo1* (*23*) and *hmgcs1* (*23*). The immune cell types identified included lymphocytes (expressing *zap70* (*32*), *il7r* (*33*) and *ifng1* (*34*)), neutrophils/monocytes (expressing *mmp13a* (*35*), *mmp9* (*36*), *lyz* (*37*) and *lect2l* (*38*)), and eosinosphils (expressing *viml* (*39*), *hp* (*39*), *cd81a* (*39*) and *gata2a* (*39*)). *Marco* (*40*), *cxcl8a* (*41*) and *csf1ra* (*42*) were highly expressed in noninflammatory macrophages, while *grn1* (*43*), *mpeg1*.*1*(*44*) and *cxcr4b* (*45*) were predominantly expressed in inflammatory macrophages. Additionally, *pax5* (*46*) and *cd79a* (*47*) exhibited peak expression specifically in B cells, while *rhag* (*48*) and transcripts of *prdx2* (*49*) and *slc4a1a* (*50*) were found only in erythrocytes (Figure 1E). Our data indicate that the cell types in zebrafish liver are conserved with those in the liver of mammals. Therefore, with the advantage of the similarity of the cell types and marker genes, we suggest that we can perform real-time tracking of their morphology and behaviors while modeling various liver diseases by marking individual liver cell types.

### Identification of hepatocyte subtypes in zebrafish

Previous studies have shown that the composition and function of hepatocytes in zebrafish are similar to those in mammals (*51*). To determine potential cell types involved in the pathogenesis of liver diseases, we performed reclustering analysis on hepatocytes and identified six distinct subclusters (Figure 2A). Our results showed that cluster 2 was the predominant type of hepatocyte (Figure 2B). To determine the characteristics of subclusters, we examined the cell proportion and DEGs in each cluster, revealing that the numbers of genes expressed were highly variable. For example, *cela1*.*5* (*52*) and *cela1*.*2* (*52*) were mainly expressed in cluster 0, while *fgg* (*53*), *cfhl4* and *apom* (*54*) exhibited peak expression in cluster 2. Moreover, *pou2f3* (*55*) and *fabp1b*.*1* (*56*) were highly expressed in cluster 4 and cluster 5, respectively (Figure 2B). The expression profiles of DEGs showed significant heterogeneity of cell subclusters (Figure 2C). To systematically investigate the features of distinct hepatocyte subtypes, we performed GO analysis for the DEGs of each cluster (Figure 2C). Our data showed that genes expressed in cluster 1 were enriched in “Oxidative phosphorylation”, “Proteasome” and “Cholesterol Biosynthesis”, and those expressed in cluster 2 were enriched in “Ribosomal Proteins”, “Response to estrogen” and “Hemostasis”. Additionally, “immune response”, “NOD-like receptor signaling pathway” and “apoptosis” were noted in cluster 3, whereas “protein processing in endoplasmic reticulum” and “cellular response to hypoxia” were noted in cluster 4. Notably, cluster 5 was enriched with genes related to “Alcohol metabolic process”, “PPAR Signaling Pathway” and “Metabolism of lipids”. Collectively, these results suggest that subpopulations of hepatocytes may play distinct roles in liver homeostasis in zebrafish. In particular, cluster 5 may be vital for lipid metabolism and synthesis. Furthermore, genes related to fatty liver diseases, such as *fabp1b*.*1, adcal* (*57*), *fabp2* (*56*), *apoa1a* (*58*), *cd36* (*59*) and *acox1* (*60*), were highly expressed in cluster 5 (Figure 2D), suggesting that cluster 5 may also be involved in the formation of fatty liver.

**Figure 2.**
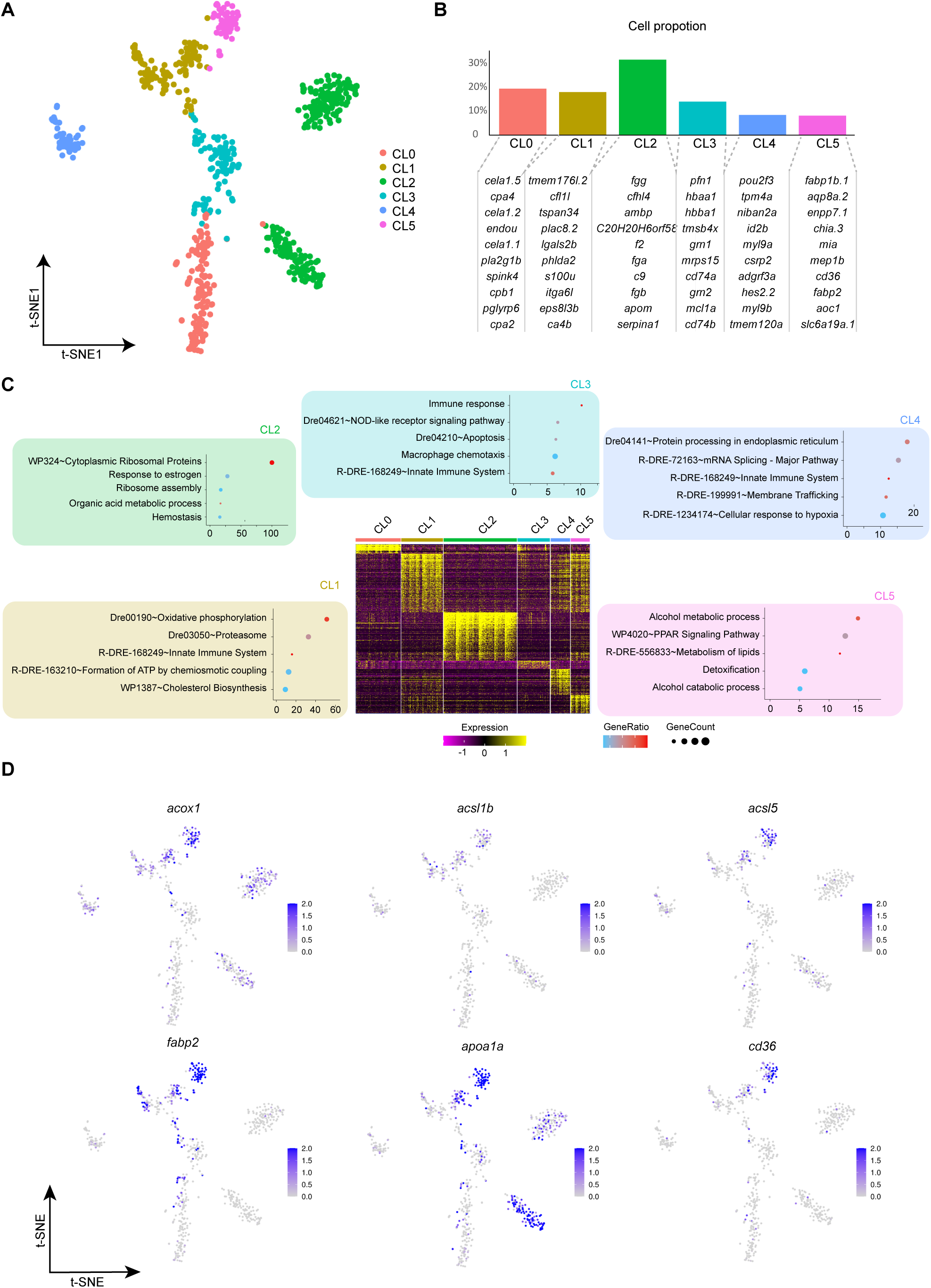
Analysis of hepatocyte subclusters. **A**. UMAP of hepatocytes reclustering. **B**. Histogram of cell proportion in hepatocyte subclusters. The upper picture represents the cell proportion of hepatocyte subclusters, and the lower picture represents the top 10 DEGs in the clusters. **C**. Bubble plot and heatmap of hepatocytes in different subclusters. The bubble plot represents the terms of subclusters in hepatocytes, and the heatmap represents the expression of DEGs in subclusters of hepatocytes. **D**. Feature plot of the key genes of various liver-associated diseases in CL5.

### The comparison of inflammatory and noninflammatory macrophages

Hepatic macrophages are key components of the liver immune system and comprise two distinct populations that possess pro- or anti-inflammatory functions (*61*). Our data showed that these two macrophage populations are present in the zebrafish liver (Figure 3A). We performed comparative analysis on the cell proportion of the two macrophage subclusters and found that there were no apparent differences (Figure 3B). Next, we performed pairwise comparisons using gene set enrichment analysis (GSEA). The functional pathways that were highly represented in the non-inflammatory macrophage population were associated with immunological tolerance, including “complement activation”, “oxidative phosphorylation”, “electron transport activity”, “fatty acid binding” and “cell–cell recognition” (Figure 3C). In contrast, the functional pathways that were highly represented in the inflammatory macrophage population were related to the inflammatory response, including the “INF-_γ_ mediated signaling pathway” and “toll like receptor signaling pathway” (Figure 3D). Furthermore, genes related to the inflammatory response, such as *ciita* (*62*), *hla-drb1* (*63*) and *tlr1* (*64*), were upregulated in inflammatory macs. To understand the underlying molecular mechanisms driving the differentiation of macrophage subclusters, we performed single-cell regulatory network inference and clustering (SCENIC). This analysis predicted the expression networks of transcription factors that were upregulated in pro- or anti-inflammatory macrophages (Figure 3E). For instance, regulons driven by members of the IRF family were enriched in inflammatory macrophages, such as *irf1*, which are essential in the development and activation of inflammatory macrophages (*65*), whereas KLF family members were enriched in non-inflammatory macrophages, such as *klf3*, which play an important role in the inhibition of inflammation (*66*).

**Figure 3.**
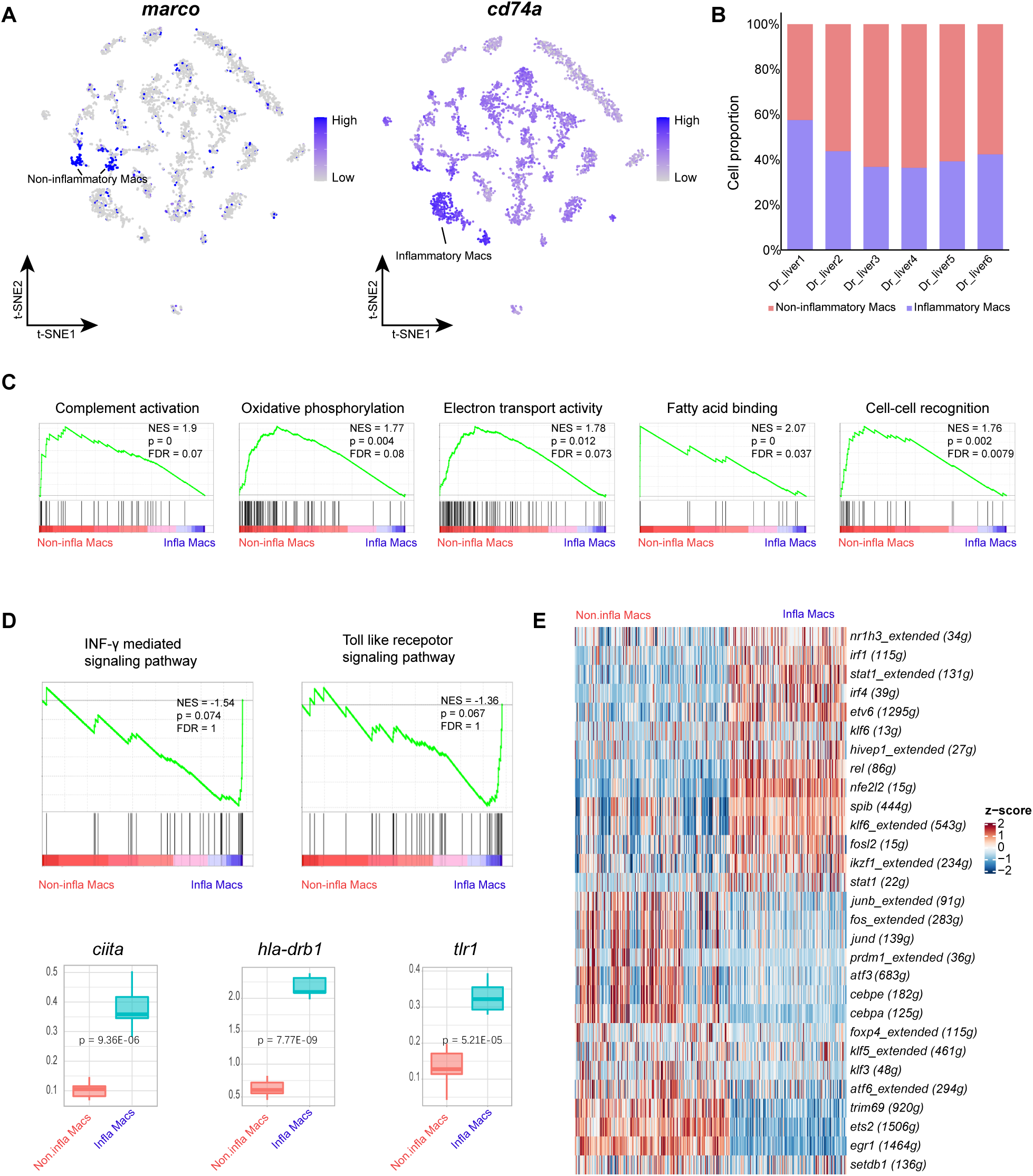
Comparison of differences between noninflammatory macrophages and inflammatory macrophages. **A**. Feature plot of the expression of key genes (marco and cd74a) in all cell types. Color scale: purple, high expression; white, low expression. **B**. Stacked histogram of the proportion of noninflammatory macrophages and inflammatory macrophages in different samples. **C**. GSEA enrichment plots of GO signaling pathways between noninflammatory macrophages and inflammatory macrophages. **D**. GSEA enrichment plots of GO signaling pathways and boxplots of key component genes associated with the inflammatory response between noninflammatory macrophages and inflammatory macrophages. **E**. Heatmap of the inferred transcription factor gene regulatory networks (SCENICs) between noninflammatory macrophages and inflammatory macrophages. Numbers between brackets indicate the (extended) regulons for respective transcription factors. Color scale: red, high expression; blue, low expression.

### Heterogeneity of T cell subclusters

The liver harbors a number of T cell populations that exert protective and pathogenic functions in infectious, inflammatory and neoplastic liver diseases (16). To gain a comprehensive view of CD4+ and CD8^+^ T cell heterogeneity in the zebrafish liver, we identified CD4^+^ and CD8^+^ T cells and further performed reclustering (Figure 4A). Based on unsupervised clustering, CD8^+^ T cells could be divided into 6 subclusters. C0_CD8-*gzmk* (*gzmk* (*67*) and *gzm3*.*2* (*67*)), C1_CD8-*rorc* (*ccl20a*.*3* (*68*), *il22* (*69*),and *rorc* (*70*)), C2_CD8-*ccl38*.*6* (*ccl38a*.*4* (*71*), *ccl38*.*6* (*71*), and *ifng1r* (*72*)) and C5_CD8-*lta* (*lta* (*73*), *tnfrsf9a* (*73*), and *egr1* (*74*)) were consistent with a memory- and/or effector-like phenotype, while C3_CD8-*mki67* (*top2a* (*75*), *hmgb2a* (*76*), and *mki67* (*77*)) and C4_CD8-*mcm4* (*mcm4* (*78*), *pcna* (*79*), and *hspd1* (*76*)) expressed markers of exhaustion and the cell cycle (Figure 4B and Figure 4D). The CD4+ T cells were grouped into 4 transcriptionally distinct states. C0_CD4-*foxp3a*(f*oxp3a* (*80*) and *tnfrsf9b* (*81*)), C1_CD4-*cd28*(*il10* (*82*), *cd28* (*83*), and *ifng1r*), and C2_CD4-*cebpb*(*cebpb* (*84*) and *eef1da* (*85*)), which were consistent with helper- and/or regulatory-like phenotypes, while C3_CD4-*mki67* (*top2a, mki67*, and *stmn1a* (*86*)) expressed markers of exhaustion and cell cycle (Figure 4C and Figure 4F). Since T cells transition to new states in metabolic diseases, monocle2 was used to reflect a possible path for differentiation of CD8+ T cell subclusters (C2_CD8-*ccl38*.*6*, C1_CD8-*rorc*, C0_CD8-*gzmk*, C5_CD8-*lta*, C4_CD8-*mcm4* and C3_CD8-*mki67* followed a continuous trajectory) and CD4+ T cell subclusters (C1_CD4-*cd28*, C2_CD4-*cebpb*, C0_CD4-*foxp3a* and C3_CD4-*mki67* followed a continuous trajectory) (Figure 4E and Figure 4G). Our analysis of finer T cell states provides better resolution of cell states and suggests testable paths for differentiation in the zebrafish liver.

**Figure 4.**
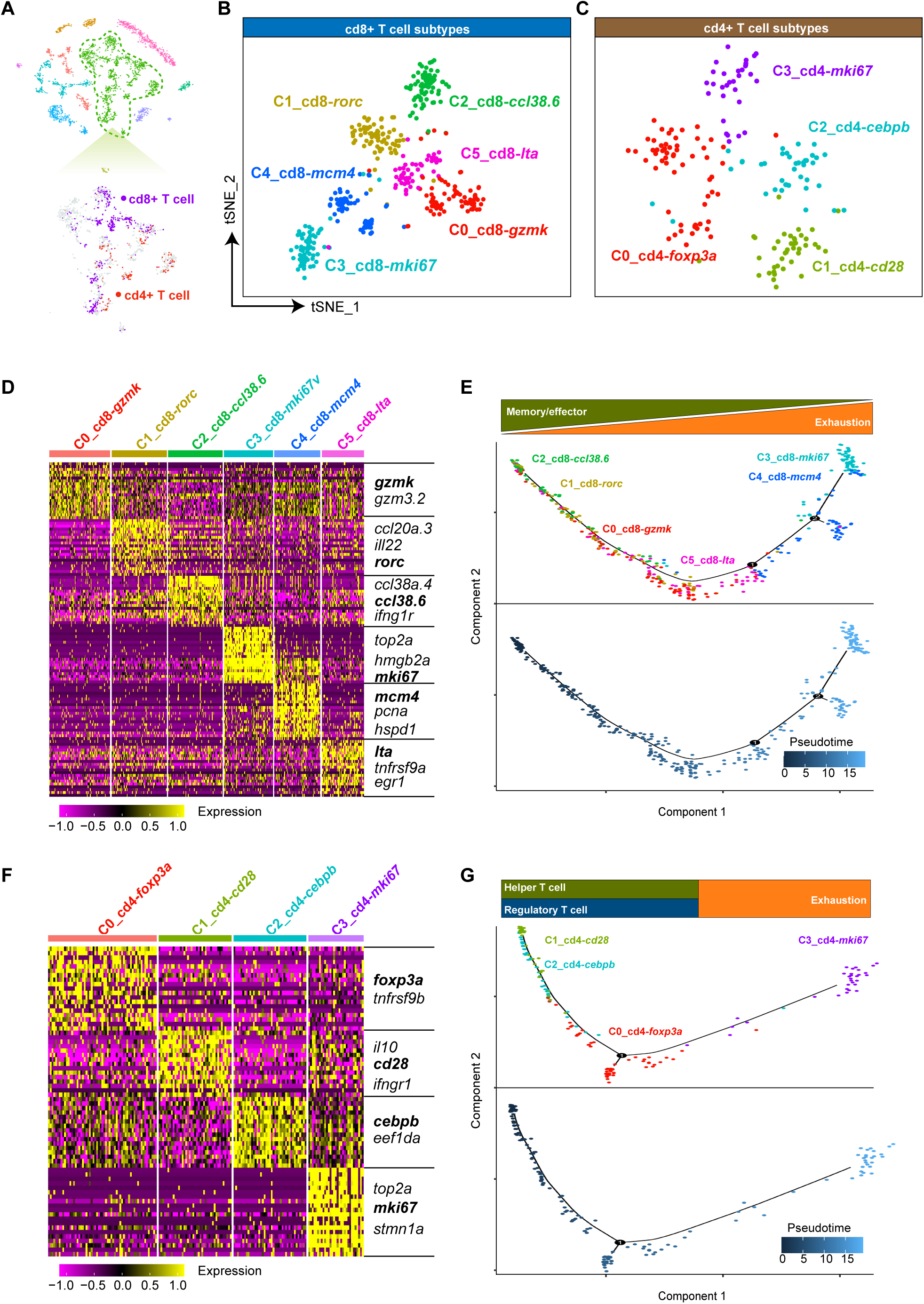
Analysis of CD8+ T cell subclusters and CD4+ T cell subclusters. **A**. t-SNE plot visualization of CD8+ T cells and CD4+ T cells, color coded for the identified T cell phenotypes. (Purple indicates CD8+ T cells, and red indicates CD4+ T cells). **B**. t-SNE plot visualization of CD8+ T cell subclusters, color coded for the identified CD8+ T cell phenotypes. **C**. t-SNE plot visualization of CD4+ T cell subclusters, color coded for the identified CD4+ T cell phenotypes. **D**. Heatmap showing the expression of the top 20 DEGs for CD8+ T cell subclusters. Z-scores were calculated by subtracting the average value for the set of data from the value for each cell and dividing by the standard deviation. **E**. Developmental pseudotime trajectory of CD8+ T cell subclusters. The upper figure represents the pseudotime process of the different CD8+ T cell subclusters, color coded for the identified CD8+ T cell phenotypes; the figure below shows the differentiation of different cell subclusters; the darker the color, the lower the degree of differentiation. **F**. Heatmap showing the expression of the top 20 DEGs for CD4+ T cell subclusters. Z-scores were calculated by subtracting the average value for the set of data from the value for each cell and dividing by the standard deviation. G. Developmental pseudotime trajectory of CD4+ T cell subclusters. The upper figure represents the pseudotime process of the different CD4+ T cell subclusters, color coded for the identified CD4+ T cell phenotypes; the figure below shows the differentiation of different cell subclusters; the darker the color is, the lower the degree of differentiation.

### Interaction between hepatocytes and immune cells

NAFLD undergoes complex pathophysiological processes, which involve a variety of factors and intrahepatic cells, including parenchymal cells and nonparenchymal cells. Importantly, these processes cannot be achieved by the actions of a single cell, rather by the interactions of different cells to form a complex cross-linking network and feedback system. However, the interaction between hepatocytes and immune cells in the liver is still unclear. Therefore, we examined potential ligand–receptor interactions between hepatocytes and immune cells to predict intercellular communication based on CellPhone DB v2.0 in normal adult zebrafish. The two macrophage subclusters identified in this study displayed relatively high numbers of interactions, while other cell types showed lower numbers of potential interactions (Figure 5A). Notably, we found that communications between hepatocytes and immune cells are bidirectional. Inflammatory macrophages were predicted to interact with Hep CL5 through the ligand/receptor pairs *ptn*/*plxn2a, cxcl11*.*6*/*dpp4, lta*/*ltbr* and *cxcl8a*/*dpp4* to promote the inflammatory state of hepatocytes, while hepatocytes regulate macrophage functions in host defense by synthesizing *mif* and *copa* combined with the expression of *cd74a* on the surface of macrophages. We also detected that *lta*/*ltbr* and *cxcl8a*/*dpp4* were active between non-inflammatory macrophages and Hep CL5 cells. Notably, the expression of *cdh1, il34* and *fam3c* by Hep CL5 could lead to the interaction with the *aEb7 complex, csf1ra* and *adgrg3*, respectively, which are expressed in non-inflammatory macrophages. These immune factors were previously reported to be associated with cell proliferation, differentiation and migration. In addition, we predicted the interaction between T cells and hepatocytes. For example, we found that the *aLb2 complex*/*f11r*.*1* showed high specificity and activity between CD8-*gzmk* and hepatocytes. *Lta* produced by CD8-*rorc*, CD8-*ccl38*.*6* and CD8-*lta* could also interact with its receptor *ltbr* in Hep CL5 cells. In contrast, Hep CL5 produced *ccl19a*, which binds to its receptor *ccr7* on CD4-*cebpb* and CD8-*gzmk* to regulate immune and inflammatory processes. *Tnfsf10*/*fas*, related to the apoptosis pathway, exhibited high activity between Hep CL5 and T cells (including CD4-*mki67*, CD8-*rorc*, CD8-*mki67* and CD8-*lta*), suggesting that hepatocytes play an important role in regulating T cell apoptosis.

**Figure 5.**
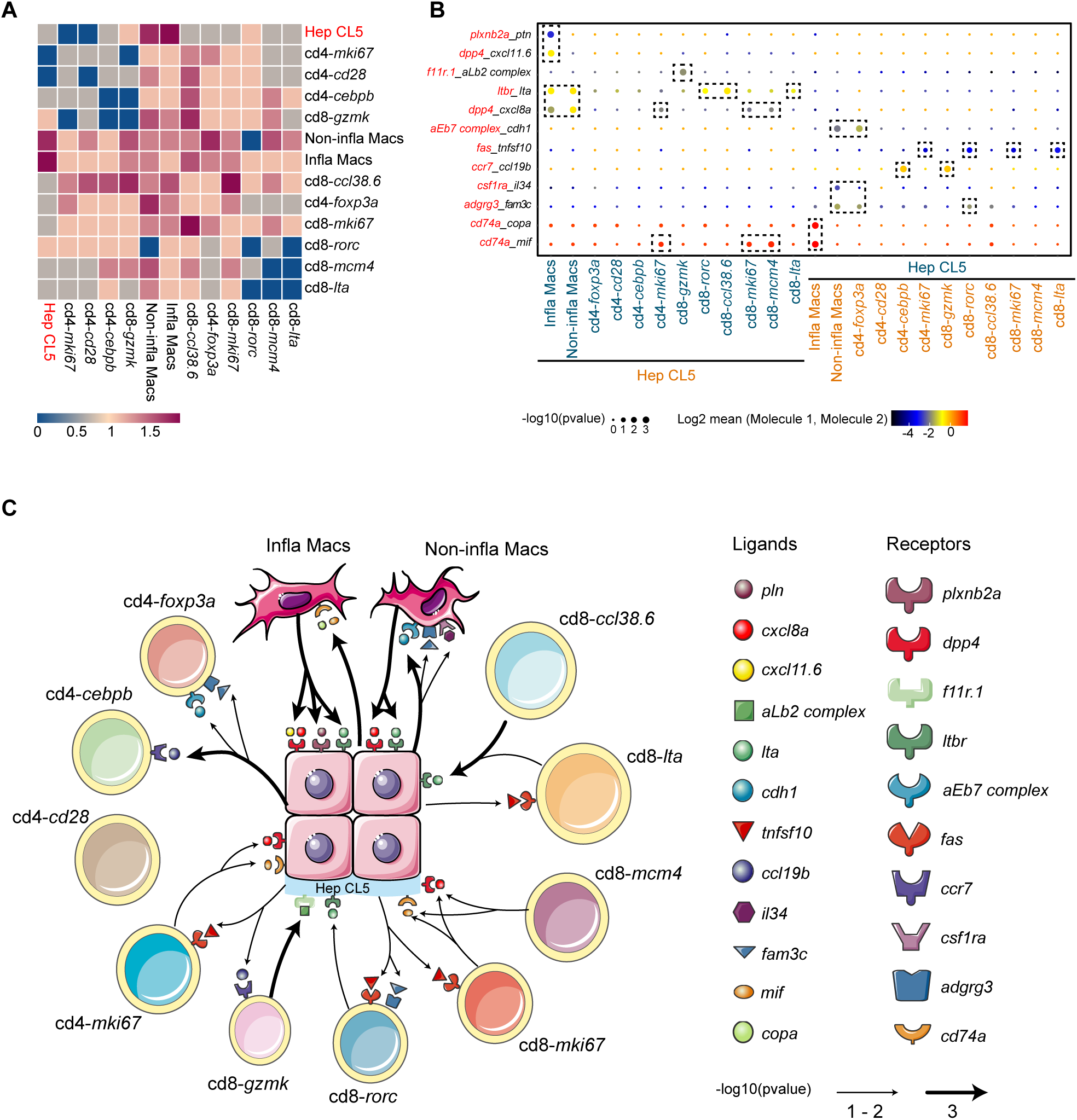
Interaction in hepatocytes-immune cells. **A**. Heatmap showing the interaction scores between hepatocyte subcluster-5 and immune cells. Color scale: Purplish red, high expression; blue, low expression. **B**. Overview of selected interactions of receptors and ligands expressed by hepatocyte subcluster-5 and immune cells. the color of the genes: red, receptors; black, ligands. **C**. Interaction pattern between hepatocytes and immune cells.

## Discussion

Zebrafish serve as an ideal animal model for the study of liver diseases. Large-scale RNA sequencing has commonly been applied to clarify the cellular repertoire in the liver. In our study, we identified 10 cell types using a single-cell sequencing approach, which is in agreement with the liver cell atlas in humans and rodents (*23, 26, 87-89*). Our data on the transcriptional profiles at the single-cell level provide novel insights into the cell repertoire in teleost organs and valuable information to explore zebrafish as an animal model to study the pathogenesis of human liver diseases.

Fatty liver is one of the common pathological symptoms of human liver diseases. It is mainly caused by the decline of liver metabolic capacity for a variety of reasons, resulting in induction of hepatocyte stress, injury and death and increased susceptibility to liver fibrosis, liver cirrhosis and hepatocellular carcinoma (*90*). Therefore, it is crucial to establish a zebrafish liver model and study the source and destination of fatty acids in hepatocytes to understand the metabolic basis of fatty liver. Notably, we found that *fabp1b1* was specifically expressed in cluster 5. *Fabp1b1* is a homologous gene of human *FABP1* and participates in intracellular lipid metabolism, such as fatty acid uptake, transport, lipid synthesis and storage (*91*) (*92*). The current study has shown that *FABP1* plays a key role in the excessive accumulation of liver fat, resulting in lipotoxicity and oxidative stress in the progression of NAFL to NASH (*93*), indicating that cluster 5 is an important group involved in lipid metabolism in zebrafish liver. In addition, we uncovered that *cd36* was also specifically expressed in cluster 5. *Cd36* is a homologous gene of human *CD36*, which is a potential biomarker for NAFLD diagnosis and patient stratification (*94*). Previous studies have shown that the cause of hepatolipotoxicity is the excessive ability of hepatocytes to manage and output fatty acids (FFAs) in the form of triglycerides (TGs). *CD36* fatty acid translocase facilitates FFA uptake and drives hepatosteatosis onset to form NASH (*94*). This suggests that cluster 5 can be used to study the role of *CD36* in lipid toxicity in the pathogenesis of NAFLD. Additionally, we also found that pathogenicity genes such as *adcal, fabp2, apoa1a* and *acox1* were highly expressed in cluster 5. In conclusion, we believe that cluster 5 could be a potential target cell model in the process of NAFLD study, and our data can provide a basis for studying the process of hepatitis in zebrafish liver.

A key finding of our study is the discovery of two distinct populations of resident macrophages in zebrafish, providing regulatory networks that affect macrophage polarization in zebrafish. Studies in humans have shown that there are two subtypes of human liver macrophages, which segregate into proinflammatory and immunoregulatory phenotypes(*23*). In our study, m*rc1b* (*95*), *hmoxla* (*96*), *marco* and *cd63* (*97*) were placed in non-inflammatory macrophages in zebrafish liver, and these genes are known marker genes for immunoregulatory macrophages. It has been suggested that zebrafish liver non-inflammatory macrophages are similar to the profiles of CD68^+^ MARCO^+^ cells in the human liver and that of resident Kupffer cells (KCs) in the rodent liver. These cells are self-renewing, nonmigratory and immunoregulatory (*23*). Conversely, the transcriptional profiles of zebrafish inflammatory macrophages resemble those of CD68^+^ MARCO^-^ cells in humans, which express high levels of *cd74a* (*45*), *cd74b* (*45*), *cxcr4b* (*45*) and *ccr9a* (*45*) and are inflammatory (*23*). In addition, it is worth noting that the *Marco* gene is a homologous gene of human *MARCO. MARCO* plays a critical role in the progression of various diseases, suggesting that the non-inflammatory macrophage cluster displaying high levels of *MARCO* expression could serve as a candidate cell type for studying the functions of liver macrophages (*23*) (*98*). It was previously reported that *IRF1* is a transcriptional regulator that is central in activating macrophages by proinflammatory signals such as the interferon-_γ_ (IFN-_γ_)-mediated signaling pathway (*65, 99*). Loss of *irf* family members *IRF1* or *IRF8* causes severe susceptibility to infections in mammals (*100, 101*). Consistent with these studies, it is not surprising that *irf1* is highly expressed in M1 phenotype macrophages (inflammatory) (Figure 2E). This suggests that they are the key transcription factors for the differentiation of M0 phenotype macrophages into M1 phenotype macrophages in zebrafish. Inflammation and fibrosis of the liver depend on the balance regulation between two KC subsets, inflammatory macrophages and noninflammatory macrophages, which are also known as M1 and M2 macrophages (*102, 103*). Previous studies revealed that during the development of NAFLD, the ratio of M1/M2 macrophages gradually increased, so the reversal of the anti-inflammatory phenotype was suggested to enhance anti-inflammatory signaling and improve the progression of NAFLD (*104*) (*2*). Hence, our study provides a further understanding of the therapeutic strategy of using zebrafish to reverse the phenotype of Kupffer cells.

Our study reveals that diverse T cells are present in zebrafish liver. We identified four CD4+ T cell subtypes and six CD8+ T cell subtypes based on T cell signature genes, such as *foxp3a, cd28, cebpb, mki67, gzmk, rorc, ccl38*.*6, mcm4* and *lta*. Zebrafish *foxp3a*, a homolog of human *FOXP3*, has proven to be a key transcription factor of Treg cells (*105*). *CD28* and *CEBPB* are involved in T cell proliferation and the differentiation of T helper CD4+ T cells, respectively (*106, 107*). A recent study has shown that GZMK+ CD8+ T cells are associated with inflammatory aging and age-related dysfunction of the immune system (*108*). In this study, we found that CD8+ T cells expressed *gzmk*, indicating that GZMK+ CD8+ T cells are present in the zebrafish liver and may play conserved roles in aging-related immune functions. *CCL3* promotes the differentiation of primed CD8+ T cells into effector cells and coordinates their migration. The *ccl38*.*6* (*ccl3* homolog) gene was expressed in zebrafish CD8+ T cells and could be important in the homing and migration of CD8+ T cells. In addition, we identified CD8-*lta*, CD8-*rorc* and exhausted T cells (i.e., CD4-*mki67*, CD8-*mki67* and CD8-*mcm4*) with high expression of cell cycle-related genes, which is consistent with observations in humans (*76*). Future investigations are required to determine whether predicted gene markers are expressed by different subtypes of T cells isolated from zebrafish.

Liver immune cells play an important role in mediating the inflammatory response and fibrosis of hepatocytes in the progression of NAFLD. We analyzed immune-hepatocyte cytokine/receptor pairs and their interaction networks and found that macrophages could be involved in regulating the inflammatory response of hepatocytes. *cxcl11*.*6*/*dpp4* showed specifically high activity between inflammatory macrophages and hepatocytes. *cxcl11*.*6* is homologous to human *cxcl9*, which was shown to be upregulated in patients with NAFLD and associated with the promotion of the progression of NAFLD to hepatocellular carcinoma (*109, 110*). It is tempting to speculate that *cxcl11*.*6* may be considered a candidate gene for studying the inflammatory response of inflammatory macrophages and hepatocytes in the context of NAFLD in a zebrafish model. *MIF* exerts antifibrotic effects in liver fibrosis via *CD74*, and the *MIF*/*CD74* interaction is considered a target for the treatment of liver diseases (*111*). Based on this, the establishment of a zebrafish *mif* knockout model is of great significance to study the progress and treatment strategy of liver fibrosis.

Our study confirmed the presence of CD74a+ hepatocytes and *mif*/*copa-*producing macrophages in zebrafish liver. Neutrophils, CD4^+^ T cells, CD8^+^ T cells, NKT cells and other immune cells are known to be involved in the pathological processes of NAFLD (*112*). Specific types of T cells, including CD8-*rorc*, CD8-*ccl38*.*6* and CD8-*lta*, were found in the zebrafish liver, suggesting that *lta*/*ltbr*-mediated interaction of CD8^+^ T cells and hepatocytes could potentially play a role in NAFLD. On the other hand, our results showed that *ccr19* secreted by hepatocytes acted on the receptor *ccr7* in CD4-*cebpb* and CD8-*gzmk* to mediate T cell migration, while *tnfsf10* secreted by hepatocytes might act on the receptor *fas* in CD4-*mki67*, CD8-*mki67*, CD8-*rorc* and CD8-*lta* to mediate the apoptosis of T cells. Our identification of these cytokine/receptor pairs in zebrafish liver provides insights into the application of zebrafish models in research into liver diseases and screening of candidate drug targets.

In summary, this work has established the first comprehensive single-cell transcriptomic atlas of zebrafish liver, which provides new insights into the molecular mechanism of zebrafish liver function. Possible candidate cells related to NAFLD and immune cell subtypes that mediate the liver immune response in zebrafish liver have been identified. Bidirectional communication between hepatocytes and immune cells and ligand/receptor interactions were detected. Our study provides a basis to explore zebrafish as a model to study liver functions and NAFLD.

## Material and methods

### Biological samples and the ethical use of animals

The experimental procedures using zebrafish were approved by the Animal Care and Use Committee at the Chinese Academy of Fishery Sciences, and all experimental procedures were performed in accordance with the guidelines for the Care and Use of Laboratory Animals at the Chinese Academy of Fishery Sciences.

### Dissociation of liver cells in zebrafish

To obtain a high-quality cell suspension, we strictly controlled the preparation time. First, the ovaries of Chinese tongue sole were dissected on ice and washed twice with DMEM, and other tissues adhered to the ovaries were removed with tweezers. The ovarian tissue was cut with a blade to form the tissue homogenate. The tissue homogenate was transferred to the combined enzyme digestion solution (0.025% trypsin + collagenase in 400μL DMEM). After heating in a 30 °C water bath for 5–10 min, digestion was rapidly terminated with 6∼7 mL of DMEM. The digested cell suspension was filtered with a 100-μm filter first and filtered again with a 40-μm filter to collect approximately 7 mL of the filtered cell suspension into a 15-mL tube. Centrifugation was performed with a horizontal rotor in a cryogenic centrifuge at 400× g for 5 min. After removing the supernatant, 7 mL of PBS was added, centrifuged at 400 × g for 5 min, and washed twice. Finally, 0.05% BSA in PBS was added, and the cells were resuspended. The cells were counted using a hemocytometer under a microscope at a concentration of 1000∼2000 cells per 1μL.

### ScRNA-seq library preparation and sequencing

The scRNA-seq library was constructed immediately after the cell suspension was prepared. We used the DNBelab C Series Single Cell RNA Library Preparation Kit based on droplet microfluidics technology for library construction. In detail, we pooled 110,000 cells per library with an average concentration of 1000 cells/μL and activity greater than 80%. Cells were prepared as droplets in which cell lysis and mRNA capture were performed using the DNBelab C4 portable single-cell system. Single-cell microdroplets were recovered by the emulsion breaking recovery system, after which magnetic bead-captured mRNA was transcribed into cDNA and subjected to 16 cycles of PCR for cDNA enrichment. Finally, cDNA products were used to prepare single-stranded DNA libraries by various steps, such as shearing, end repair, ligation, 12 cycles of PCR, denaturation, circularization, and digestion. Then, 10 ng of the digested product was obtained for sequencing using the MGISEQ 2000 platform.

### ScRNA-seq data preprocessing

The raw data obtained by scRNA-seq using the MGISEQ 2000 platform were filtered and demultiplexed using PISA (version 1.10.2) (https://github.com/shiquan/PISA). Reads were aligned to the reference genome from NCBI using STAR (version 2.7.9a) and sorted using sambamba (version 0.7.0). The cell versus gene UMI count matrix was generated using PISA.

### Quality control, normalization, removal of doublets and dimensionality reduction

For every sample, we used Seurat (https://satijalab.org/seurat/) to obtain ideal cells defined as those expressing the number of genes detected between 200 and 3500, with a mitochondrial percentage of less than 5%. Using the filtered cells, we performed normalization by the SCTransform function and followed the official website tutorial to remove doublets using the R package DoubletFinder. Subsequently, for visualization, we performed dimensionality reduction of the data using RunPCA, and the first 50 principal components (PCs) were computed with the 3000 highly variable genes using log-normalized expression values calculated by the SCTransform function. Using the ElbowPlot function, we chose the first 30 PCs for the next steps, including computing the k.param nearest neighbors by the FindNeighbors function, constructing the SNN graph by the FindClusters function, and computing the t-SNE by the RunTSNE function. We saved rds to conduct further downstream integration.

### Integration, cell cluster and DE analysis

Using the SelectIntegrationFeatures function, we retained the top 3000 scoring features to use when integrating all samples, and we found the anchor features by the FindIntegrationAnchors function, with the normalization method parameter as “SCT”. Then, we performed integration by the IntegrateData function. Subsequently, we repeated the above method to recluster the cells. Finally, we performed differential expression (DE) using the FindAllMakers function.

### Gene pathway analysis

Gene function enrichment analysis was performed using Metascape (https://metascape.org/gp/index.html).

### GSEA Analysis

To identify the significantly enriched pathways in the difference of noninflammatory mac and inflammatory mac, we used Gene Set Enrichment Analysis (GSEA, http://software.broadinsitute.org/gsea/index.jsp) to perform enrichment analysis. We first transformed the zebrafish single-cell data matrix into an adult matrix. The 8831 one-to-one homologous genes of humans and zebrafish were obtained by BLAST.

### SCENIC Analysis

To perform transcription factor network inference, we used the SCEINC R package (version 1.2.4, which corresponds to RcisTarget 1.8.0 and AUCell 1.12.0) to calculate the activity of the regulatory networks on the inflammatory mac and noninflammatory mac datasets using default settings.

### Cell–cell communication analysis

We extracted CD4+ T cell subclusters, CD8+ T cell subclusters, noninflammatory macrophages, inflammatory macrophages and hepatocyte cluster 5 from the transformed matrix as input and then used the CellPhoneDB Python package to perform cell–cell communication analysis. The algorithm is based on the receptor expression of one cell type and the corresponding ligand expression of another cell type to deduce the enriched receptor ligand interaction between the two cell types. Then, we identified the most relevant cell type-specific interactions between ligands and receptors and only considered the receptors and ligands highly expressed in the corresponding subpopulations.

## Acknowledgements

This work was supported by the National Key R&D Program of China (2018YFD0900301); the AoShan Talents Cultivation Program Supported by Qingdao National Laboratory for Marine Science and Technology (2017ASTCP-ES06); the Taishan Scholar Project Fund of Shandong of China to C.S.; the National Ten-Thousands Talents Special Support Program to C.S.; the Central Public-interest Scientific Institution Basal Research Fund, CAFS (No.2020TD19).

## Author contributions

Changwei Shao designed and supervised the research. Xiang Liu, Jian-Yang Chen, and Xianghui Zhang performed all experiments, collected and interpreted the data. Yingyi Huang and Yifang Lu did the bioinformatics analysis of single-cell transcriptomic data. Hong-Yan Wang, Yubang Li, Zhongdian Dong, Kaiqiang Liu, Zhongduo Wang, Qian Wang, Guangyi Fan, Jun Zou, and Shanshan Liu provided essential reagents and suggestions. Xiang Liu and Hong-Yan Wang collected the fish samples. Yingyi Huang and Xiang Liu wrote the manuscript. Yingyi Huang, Xiang Liu and Changwei Shao edited the manuscript and revised it critically. All authors took part in the interpretation of data.

## Figure legends

**Dataset supplement 1 DEGs in all cell types in zebrafish liver**.

**Dataset supplement 2 DEGs in all cell types in hepatocyte subclusters**.

**Dataset supplement 3 DEGs in all cell types in CD4+ T cell subclusters**.

**Dataset supplement 4 DEGs in all cell types in CD8+ T cell subclusters**.

